# Dynamical predictors of an imminent phenotypic switch in bacteria

**DOI:** 10.1101/145821

**Authors:** Huijing Wang, J. Christian J. Ray

## Abstract

Single cells can stochastically switch across thresholds imposed by regulatory networks. Such thresholds can act as a tipping point, drastically changing global phenotypic states. In ecology and economics, imminent transitions across such tipping points can be predicted using dynamical early warning indicators. A typical example is "flickering" of a fast variable, predicting a longer-lasting switch from a low to a high state or vice versa. Considering the different timescales between metabolite and protein fluctuations in bacteria, we hypothesized that metabolic early warning indicators predict imminent transitions across a network threshold caused by enzyme saturation. We used stochastic simulations to determine if flickering predicts phenotypic transitions, accounting for a variety of molecular physiological parameters, including enzyme affinity, burstiness of enzyme gene expression, homeostatic feedback, and rates of metabolic precursor influx. In most cases, we found that metabolic flickering rates are robustly peaked near the enzyme saturation threshold. The degree of fluctuation was amplified by product inhibition of the enzyme. We conclude that sensitivity to flickering in fast variables may be a possible natural or synthetic strategy to prepare physiological states for an imminent transition.

## 1. Introduction

Regulatory networks are subject to biochemical noise, and have nonlinearities such as ultrasensitive thresholds embedded within [1]. These properties make predicting single-cell phenotypic transitions challenging [2]. Thresholds in the network often demarcate the location in phase space of critical transitions, responsible for producing discrete phenotypes [3]. For example, bacterial molecular regulatory networks contain ultrasensitive thresholds at stoichiometric points of strong protein-protein or protein-RNA interactions, and metabolic networks have critical thresholds at the point of enzyme saturation [4–6].

When a network is poised near such a threshold, fluctuations from intrinsic noise can result in fractions of the cellular population being on either side of the threshold. This may lead to so-called bet-hedging, where subpopulations are pre-disposed to survive in different unpredictable future environments [7]. A medically urgent type of this is in the formation of the persister phenotype: a slow-growing, antibiotic-tolerant subpopulation [8]. Here, network thresholds create a probability of fast-growing cells to enter persistence either intrinsically or induced by an extrinsic stress [9]. We previously identified a novel mechanism for persister formation by crossing a phenotypic threshold apparently determined by enzyme saturation [5]. In conditions that increase the probability of crossing the growth arrest threshold, the fraction of cells that are persisters is enriched. Metabolic changes also underlie the formation of cancer [10], suggesting that metabolic thresholds may result in phenotypic shifts within tumors.

In ecological and economic studies, critical transitions in complex networks have characteristic dynamical signatures that are capable of predicting them, such as "flickering" dynamics, where a fast-changing variable rapidly fluctuates between two states from intrinsic noise when the system is poised on the threshold between them. Methods have been developed for analyzing flickering to predict critical transitions [11–15]. Early warning indicators have been used to predict disease progression as well [16, 17]. Analogously, fluctuations occurring in cellular regulatory networks near a threshold, such as enzyme saturation, may originate from heterochronicity of the fast processes of post-translational dynamics (metabolic fluctuation) and the slow process of protein synthesis (figure 1).

**Figure 1.**
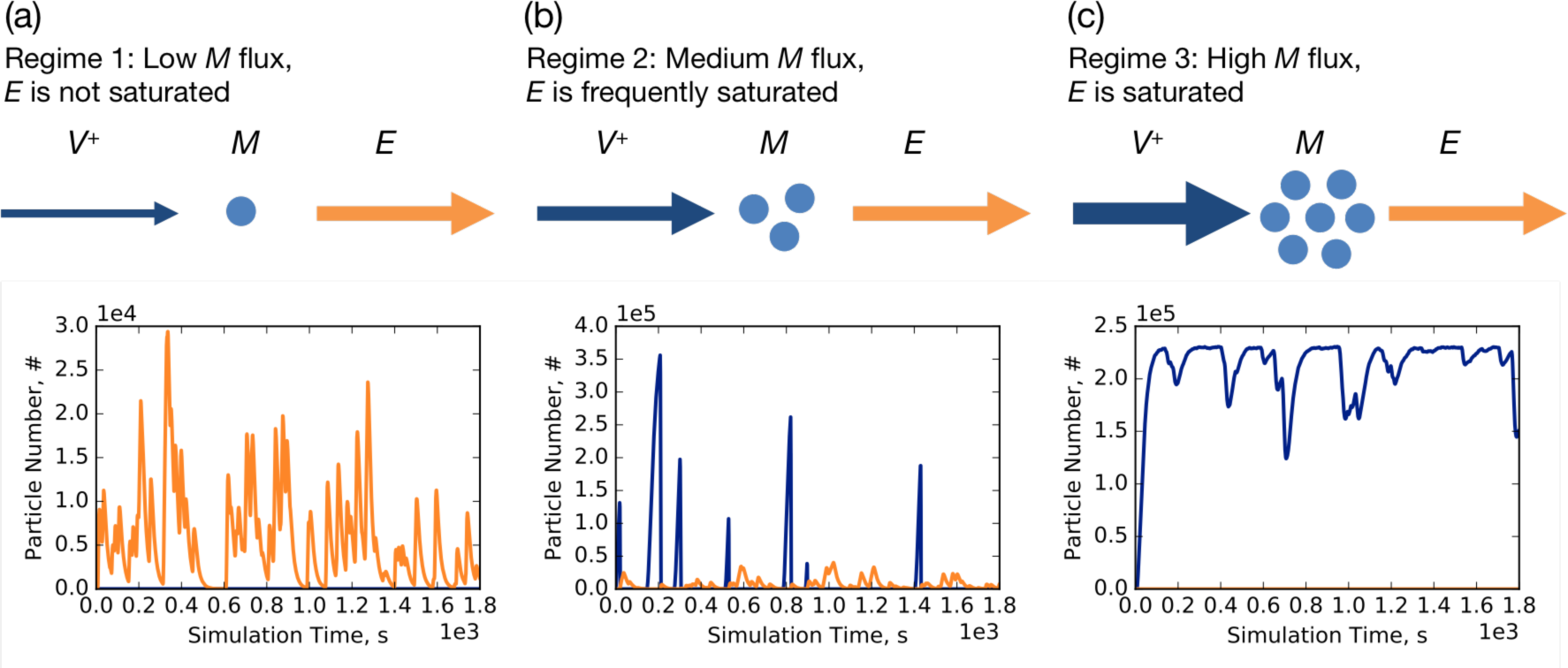
Cartoon representation of three dynamical regimes of a bacterial metabolic step with typical time series from corresponding stochastic simulations. Blue arrows represent metabolic precursor influx *V*^+^, and orange arrows represent enzyme-mediated consumption. The lower panels show typical time series. Orange lines show free enzyme time trajectories and blue lines show metabolic precursor time trajectories. (a) Regime 1 is an unsaturated system with low influx *V*^+^ relative to enzyme level. (b) Regime 2 is the expected flickering regime with random oscillation between saturated and unsaturated system states with intermediate influx *V*^+^ near the critical threshold. We observe that metabolite spikes appear when the enzyme in the system is saturated. (c) Regime 3 is a saturated metabolic step with high influx *V*^+^. Here the metabolic precursor *M* fluctuates around a high steady state, and enzyme is always saturated.

In reproducible conditions, flickering may be an indicator of near-criticality that governs phenotypic transitions. It may also be useful for identifying the dynamics of cells driven by variables moving much more slowly than molecular fluctuations. Therefore, it could be developed as a tool for characterizing intact molecular networks.

Motivated by these observations, we asked how predictable the crossing of a metabolic phenotypic threshold is at the level of a single cell. Various studies have explored the parameter space that allows the saturation threshold to have an effect on metabolite dynamics [4, 18]. We explored dynamical regimes near the enzyme saturation threshold in a simple metabolic step, and compared how different factors can influence system behaviors. Using stochastic simulations and analytical noise estimates of the stationary system, we determined the roles of enzyme affinity, burstiness of enzyme gene expression, product inhibition, and rates of metabolic precursor influx. We evaluated metabolite dynamics as early warning indicators under the assumption that changing external conditions are slow enough to permit detection of metabolite dynamical states, and used a quasi-steady state assumption for the external driver of the phenotypic switch (here, the metabolite influx rate).

We conclude that spiking rate and spike span time of metabolic precursor in a single cell can work as dynamic system indicators in a reasonable parameter space of the model, suggesting that studies predicting phenotypic transitions in larger networks are a promising future direction.

## 2. Methods

### 2.1. Linear Noise Approximation of a Minimalistic model

We analytically solved a minimal 1-variable model and used *Mathematica 11.0* to numerically integrate differential equations. We used the linear noise approximation (LNA) to predict noise levels (section 3.1) by solving the stationary fluctuation matrix equation as shown previously [19, 20]. Potential analysis for steady state metabolic precursor followed Strogatz [21]. Detailed results are given in the model supplement Mathematica notebook.

### 2.2. Stochastic Simulations

Detailed models followed the reaction scheme in figure 2a. The models were first tested in COPASI [22] to verify functionality and to pre-evaluate parameter ranges. For large scale simulations, we converted the verified model into the *StochKit* [23] file format, and submitted simulation jobs to University of Kansas Advanced Computing Facility (KU ACF) cluster. All model file handling was done with Python 2.7.9. For each set of parameters, we ran simulations for 180000 seconds of simulation time with 100 or more repeats using the τ-leaping method in *StochKit.* Simulated data were sampled every 1 second. We chose random seed 1 for each set of simulations to allow repeatability of results. The detailed reaction scheme and parameters are shown in tables S1-S5. Example simulation files are given in the model supplement.

**Figure 2.**
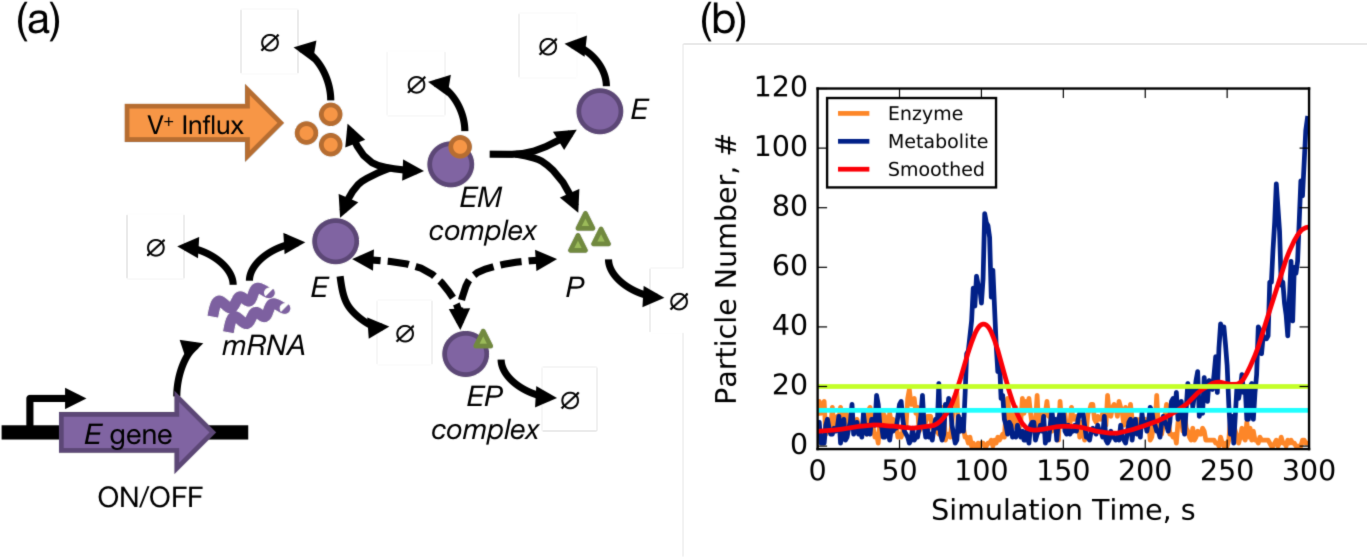
Characterizing flickering in stochastic simulations. (a) Detailed simulation scheme, including enzyme *E* gene transcription and enzyme-metabolite interactions *E: M* or *E*: *P*. Dashed arrows indicate negative feedback reactions by metabolic product *P* that competitively inhibits *E:M* interactions. (b) Characterizing metabolic spikes. Simulation trajectories can be noisy and smoothing helps retain spike information while filtering out smaller fluctuations. In this figure, *M* time trajectory is shown in blue, smoothed *M* trajectory is shown in red, and *E* trajectory is shown in orange. Free enzyme threshold *θ_e_ = 20* is indicated by the light green bar, and the metabolic precursor threshold *θ_Μ_* is the cyan bar. The intersection of cyan bar and red line characterizes the spikes, where spike span time *T_sp_*” is the *x*-axis distance between intersections. Two intersections define one spike.

### 2.3. Detecting and Quantifying Flickering in Time Series Data

In complex systems, including ecological systems and cellular regulatory networks, the dynamics can be noisy with nonlinear sensitivities amplifying the noise [2, 16, 24]. Though the flickering we quantify here has essential differences from bistable flickering previously observed in ecological transitions, we will find that the common behaviors of rising noise near threshold, and loss of resilience, result in a corresponding increase in spike rates and spike span time near the ultrasensitive threshold of a metabolic step, in common with bistable critical thresholds.

We characterized flickering of metabolite in our stochastic model (figure 2a) using a three-step data processing procedure after obtaining simulation time series (figure 2b). First, we defined an enzyme threshold to establish enzyme saturation states. We empirically determined *θ_e_ = 20* for our chosen parameter set. Thus we define that, for a time trajectory at given time *t,* when free enzyme *E*(t) ≥ *θ*_e_, enzyme in the system is not saturated, while if free enzyme *E*(t) < *θ*_e_, the system is in a saturated state. By collecting the time point *T_sa_* when the system is in the saturated state, divided by total simulation time *T_tot_*, we have the percentage of enzyme saturation for one simulation time trajectory:

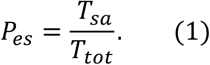

We also collected the maximum value *M_max_* of corresponding metabolic precursor quantity *M(t)* when the system is in the unsaturated state, and set *θ_Μ_ = M_max_* + 3 as our threshold for identifying spikes. Second, we performed simulation time series smoothing to filter out non-significant noise and rapid regime drifts, while preserving enough spike information. A Gaussian filter [25] is applied for this step to smooth metabolic precursor time series *M(t),* with window size of 39 data points and standard deviation of 10 molecules/cell. Third, we obtained our smoothed data set of metabolic precursor *M_post_*(t), we compared this data set with the metabolic precursor threshold *θ*_Μ_, and thus identified spikes as shown in figure 2b. The intersections of *M_post_ (t)* with *θ_Μ_* are regarded as the base points of the spikes. Here we count the spike number *N_s_* as:

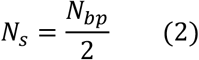

Where *N_bp_* is the number of spike base points. Thus, spike rate *β_sr_.* is defined as:

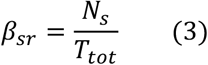

We also collected the spike span time in simulation trajectories, a quantity related to system resilience. System resilience refers to its capacity to absorb changes and remain in similar states and structure [26]: when a system is approaching a threshold, loss of resilience accompanies increasing return times in many studies [11, 27]. Spike span time *T_sp_* indicates the time needed for a temporally saturated system to return to unsaturated state, which is the distance between intersection points of smoothed time series *M_post_ (t)* and threshold *θ_Μ_* for one spike. For each set of parameter sweeps, we simulated 100 repeats and calculated the averages 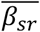 and 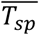, and mapped their values to heatmaps corresponding to their parameter space. Example analysis Python files are given in the model supplement.

### 2.4. Comparison of Established Indicators with Spike Rates and Spike Span Time

#### 2.4.1 Comparing Early Warning Signals

We compared spike rate and spike span time to various established indicators using the "earlywarnings" toolbox [14] for R [28]. Similar to the ecological and economic time series where an external driver drives the system across a critical threshold, we simulated a time trajectory with increasing values of *V*^+^ over time. Using the regulatory network shown in figure 2a, we added a time trigger to the model that changed parameter *V*^+^ every 90,000 seconds from 0 to 1000 /s with 20 jumps, and simulated the model for 10^6^ seconds. The long simulation time for each parameter value step is intended to allow enough time for the system to reach a quasi-steady state at each step. The simulation produces rather large amounts of data for the R toolbox to calculate in a reasonable time, so we lowered the sampling resolution using 4000 evenly spaced intervals. We tested the performance of a few commonly used early warning signals along with spike rate and spike span time.

#### 2.4.2 Window size and white noise related sensitivity

The performance of early warning indicators depends on characteristics of data and the data preprocessing details. Here we consider the noise and data pre-processing window size effects on the robustness of the indicators’ performance. Based on the results from early warning comparison (below), we tested the robustness of spike rates, spike span time, autocorrelation-at-lag-1 (ar(1)), and standard deviation (SD), based on performance or the fact that they are commonly used indicators in the case of ar(1) and SD.

To test the robustness of indicators regarding window size, the same trajectory data as the early warning comparison was used, and we varied both rolling window size for indicator calculation and filtering window size for data detrending and smoothing, and plotted the indicator time series. The window size *η* ϵ [0,1] is interpreted as the percentage of the total length of the time series data, and the filter window size *ζ* is the number of data points in each filter window. For testing robustness of ar(1), we changed rolling window *η* from 1% to 31% of the trajectory time length, and filtering window size *ζ* from 1 to 22 data points. For testing robustness of spike rates and spike span time, rolling window *η* changes from 1% to 52% of the trajectory time length, and filtering window size *ζ* changes from 1 to 100 data points.

To test robustness to extraneous noise, we added a scaled noise *σ* term to the trajectory as follows:

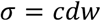

where *c* is the noise scale factor, and *dw* is standard normally distributed noise. Parameter *c* is assigned one of 5 values: 10^3^, 10^4^, 10^5^, 10^6^, 10^7^. After adding scaled noise, we then performed indicator calculations to assess their performance.

#### 2.4.3 Indicator significance and test model

We tested the statistical significance of the indicator to determine true critical transitions and avoid false positive indicators (type I error), and to use as a testing method for applying spike rate and spike span time as early warning indicators the system crossing a critical transition. For this purpose, we used a two-sample Z-test applied to spike rate and spike span time.

We used the BDS test [29] to detect nonlinear serial dependence in time series data. With properly detrended data, BDS tests the null hypothesis *H_0_* that remaining residuals are independent and identically distributed (i.i.d.). Rejecting the BDS null hypothesis *H_0_* suggests nonlinearity or nonstationarity in the time series.

### 2.5 Enzyme Gene Burstiness Definition and Related Deductions

Parameter *P*_on_ is the probability of the enzyme gene (single copy) being in the ON state, and can be described as following:

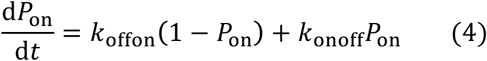

By solving the steady state, we get

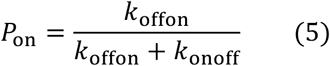

For mRNA_fi_, we solve the steady state and find

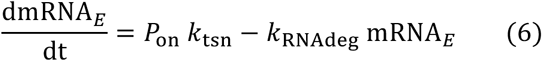

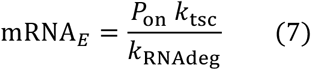

To keep enzyme expression levels constant, and then compare the effect of burstiness, we have a constant steady state enzyme number in the system:

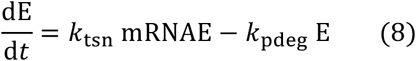

Solving for enzyme steady state, we find

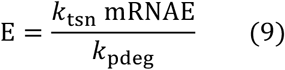

Substituting steady state expressions (5) and (7) into (9), we have

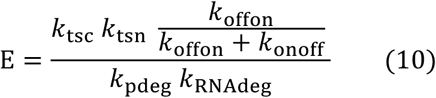

For a fixed *P*_on_, we can calculate *k*_onoff_ by

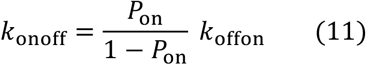

### 2.6 Simulation Trajectory Phase Indicating Phenotypic Changes in Single Cells

In our study, different phenotypes refer to distinct metabolic precursor phases in single cells. We used the percentage of saturated enzymes to indicate simulated system states, due to their closely related dynamic behaviors. We defined the system state with the following procedure. First, we collected the final time point data *M* for all repeated trajectories, and the average 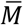 was calculated and regarded as the expected steady state value. Then we set thresholds to define three system states (phenotypes) at the final time: the low metabolic precursor state, the transition state, and the high metabolic precursor state. As the sensitivity of a threshold is shown by the EC90: EC10 ratio (EC - effective concentration of signal output [30]), we take EC10 and EC90 as thresholds and for trajectory *i* the system state Σ_*i*_ is defined as following:

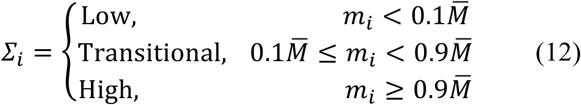

Finally, the number of trajectories in different states is obtained for each parameter sweep simulation (the distribution of three states in each set of simulation is shown in Fig. S1, S2 and S3). Here we observe that in all cases, near the enzyme saturation threshold, the system is likely to be in a bimodal distribution, corresponding to a phenotypic switch.

## 3. Results

To determine the generality of dynamical markers near a critical transition in metabolism, we first used a minimalistic model with a single metabolite variable subject to degradation by a Michaelis-Menten process. We then studied various dynamical consequences of gene burstiness, enzyme gene transcription, enzyme affinity, and metabolic product inhibition using stochastic simulations in a more detailed model.

## 3.1 Noise Regimes in a Minimal Metabolic Step Model

In a metabolic network containing an ultrasensitive threshold from enzyme saturation, there are three distinct regimes, as shown in figure 1. Different weights of blue arrows indicate different metabolic precursor *M* influx rates, *V*^+^, and metabolite consumption is indicated by orange arrows. Regime 1 (figure 1a) corresponds to low metabolic precursor influx *V*^+^, where the metabolic precursor *M* concentration is low when enzymes are not saturated. Regime 2 (figure 1b) corresponds to moderate metabolic precursor influx *V*^+^ that pushes the system near the enzyme saturation threshold, where metabolic precursor *M* concentration is frequently fluctuating, thus forming spikes (figure 1b blue line). Regime 3 (figure 1c) corresponds to high metabolic precursor influx *V*^+^, pushing the system across the enzyme saturation threshold to achieve a high *M* concentration.

To explore fluctuations in the three regimes shown in figure 1, we wrote a minimal model having constant metabolic precursor *M* influx with rate *V*^+^, enzyme *E* consumption approximated by Michaelis-Menten kinetics, and dilution from cell growth:

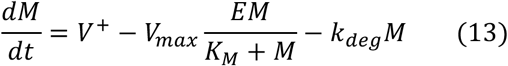

where *K_M_* is the classic Michaelis-Menten constant, *V_max_* is the maximal turnover rate of the enzyme *E*, and *k_deg_* is the dilution rate of metabolic precursor *M*. We analytically solved the differential equation; the regimes of metabolic precursor *M* are shown in figure 3a. The model predicts three qualitatively distinct metabolic states in the system and an ultrasensitive switch. At low *V*^+^, *M* has a low steady state, and after crossing the threshold, *M* has a high steady state. The red dashed line in figure 3 indicates the threshold. A slight perturbation near the enzyme saturation threshold with varied *V^+^* can induce dramatic changes in metabolic precursor *M* state change in the system.

**Figure 3.**
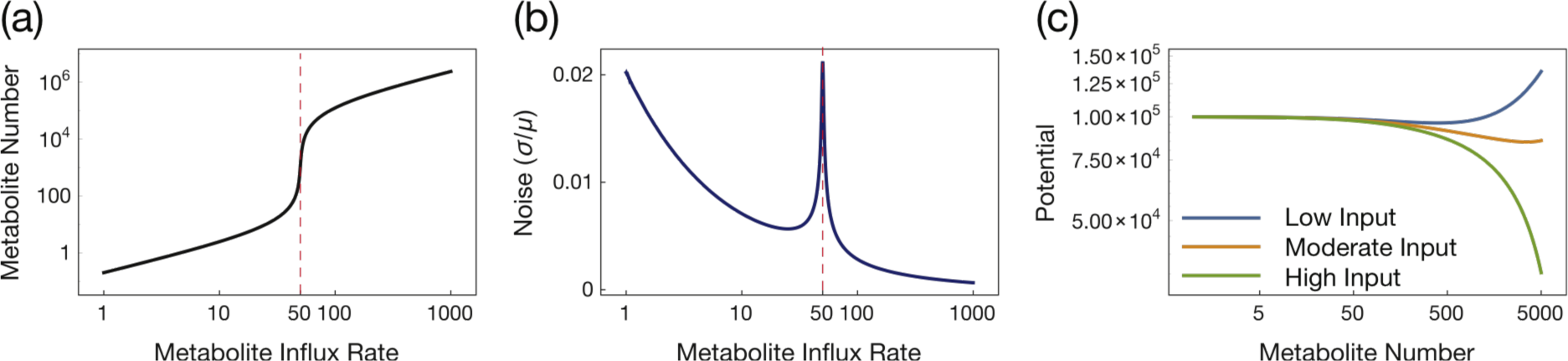
Analysis shows distinct dynamic regimes with a minimalistic model. The model is described in the main text. (a) By solving for the differential equation steady state, we obtained the phase graph of metabolic precursor *M* quantity changing with varied *V*^+^. Parameters were chosen as follows: *E =* 50, *K_M_ =* 10, *k_deg_ =* 0.004, *V_max_ =* 1. (b) The linear noise approximation (LNA) result shows that noise of the system peaks near the threshold, consistent with the observation that flickering occurs near the threshold. Noise is plotted as coefficient of variation (σ/μ). (c) Steady state *M* potential *U* under different *V^+^* conditions is calculated with the method described in the main text. Three distinct potential landscapes show that small perturbations of *V^+^* near the threshold can significantly impact system dynamics, and flickering forms from the random switches between those regimes. Parameters were chosen as follows: low input: *V^+^ =* 40, medium input: *V^+^ =* 50, high input: *V^+^ =* 60.

To approximate noise levels in the system, we used the LNA. Clearly, metabolic noise peaks at the enzyme saturation threshold marked by the red dashed line shown in figure 3b, predicting that flickering most probably appears near the ultrasensitive threshold. We note that overall noise levels are low in this simple, single-variable model. As we will show below, gene expression dynamics can enhance fluctuations around the threshold.

Notably, flickering here is different from ecological studies, where it emerges from a shift between two steady states [27]. In our minimal model, there is only one steady state for each set of parameters (varied influx *V*^+^) in the system. Potential analysis for the steady state demonstrates distinct potential landscapes for the three regimes (figure 3c). For a differential equation system in the form of:

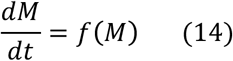

The potential is defined as:

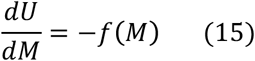

where *U* represents the potential [21]. We solved
Equation (15) for the steady state metabolic precursor *M,* and chose three *V*^+^ values near the ultrasensitive threshold. As in figure 3c, *V^+^ =* 40 corresponds to regime 1, where the enzymes are not saturated and the system possess a low *M* steady state, shown as a potential well with blue curve. *V^+^ =* 50 corresponds to regime 2, where the system is near the enzyme saturation threshold, shown as a relatively flat curve in orange. *V^+^ =* 60 corresponds to regime 3, shown as a green fast descending potential decline, implying a system preference for shifting to a high metabolite concentration. We will show that flickering emerges in metabolic steps from noise driven shifts across a nearly flat potential landscape at intermediate input.

## 3.2 A Detailed Model to Characterize Flickering Regimes

Based on the minimal model, we developed a more detailed metabolic step model where reactions are described by mass action laws. As shown in figure 2a, we integrated enzyme gene regulation, transcription and translation, enzyme dynamics, metabolic precursor influx, and product inhibition (feedback) into the model. Using this model, we tested the hypothesis of flickering as an ultrasensitive threshold predictor, and explored parameter space to understand how general the phenomenon is.

### 3.2.1 Comparison of Indicators Reveals Reliable Detection of Metabolic Flickering

To determine useful indicators for metabolic flickering, we did a comparison of various indicators that have been used previously along with spike rate and spike span time. We generated a trajectory with gradually increasing influx of metabolite over time. We used detrended data of modeled metabolite to calculate the following indicators: spike rate, spike span time, autoregression, autocorrelation, return rate, density ratio, standard deviation, and skewness (see section 2.3). We additionally computed non-parametric drift-diffusion-jump metrics for the time series: conditional variance, total variance, diffusion, and jump intensity.

Different early warning indicators have various responses to the critical transition, as shown in figure 4. We used filter window size *ζ =* 39 for data smoothing and detrending, and window size *η =* 10% of the time trajectory to calculate indicators. According to figure 4, autoregressive-coefficient-at-lag-1 (ar(1)), autocorrelation-at-lag-1 (acf(1)), return rate, density ratio, and standard deviation exhibit abrupt changes in trend when crossing the critical transition, while spike rate, spike span time, and skewness show a peak near the transition. The density ratio exhibits an increasing but oscillating pattern, which is not a reliable indicator for the transition in this case. We also calculated nonparametric drift-diffusion- jump metrics for the simulated data, as shown in the figure S4. This analysis shows large fluctuations near the transition, but with too much noise to use as an early warning indicator. We prefer spike rate and spike span time due to its direct implication of metabolite flickering and enzyme saturation. Spike rates and spike span times could also be captured in microscopy experiments if there is an appropriate metabolic probe.

**Figure 4.**
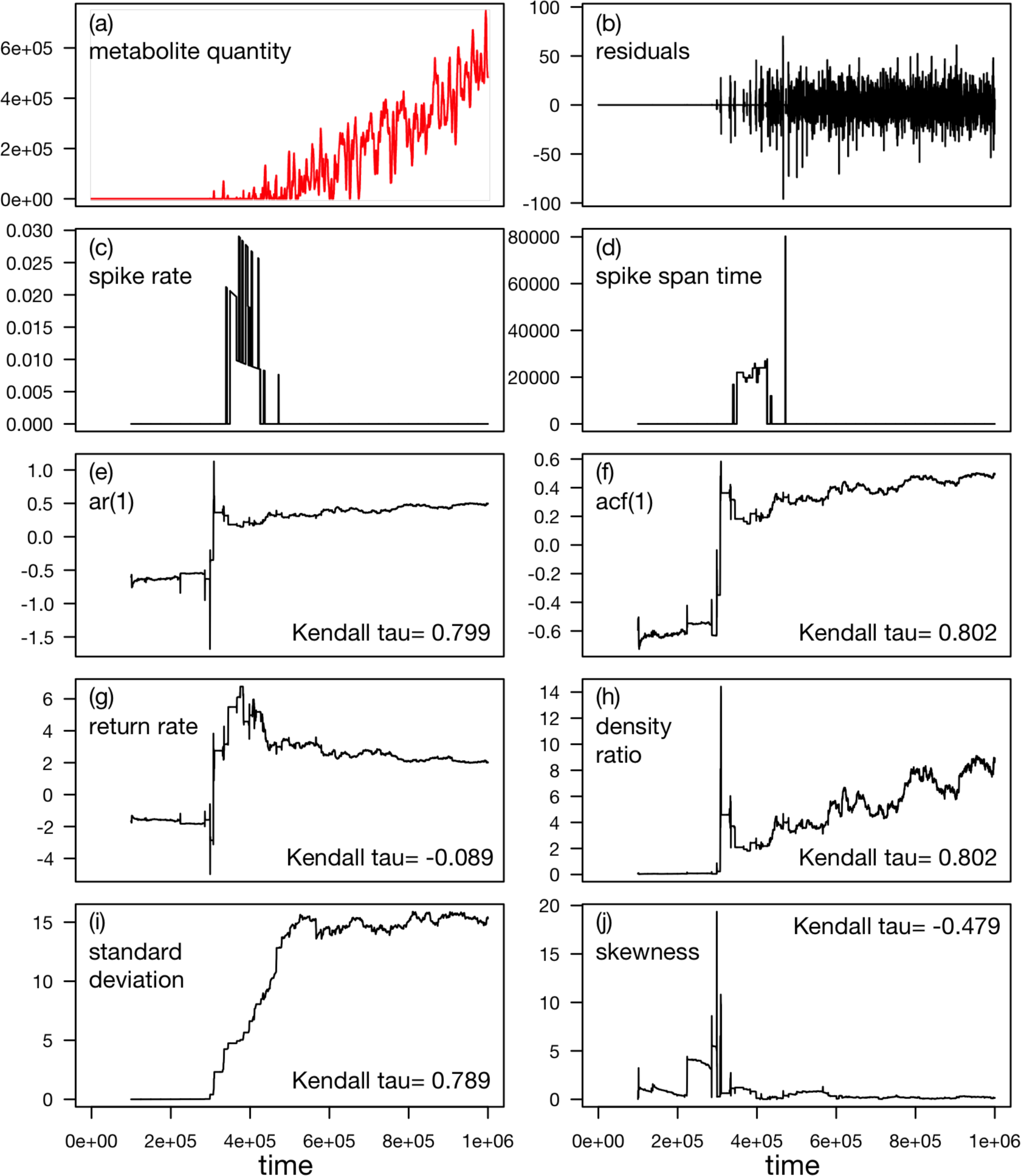
Comparison of early warning indicators for simulated metabolite data. (a) Time series of simulated metabolite quantity after filtering. We simulated the network described in figure 2a for 10^6^ seconds with *V^+^* increasing over time as described in section 2.4. (b) Residual time series after applying Gaussian filter with filter window size of 39s. (c - j) Early warning indicators of spike rate, spike span time, autoregressive-coefficient-at-lag-1 (ar(1)), autocorrelation-at-lag-1 (acf(1)), return rate, density ratio, standard deviation, and skewness calculated with window size of 10% of the trajectory. Except for spike rates and spike span time, all EWSs are defined and calculated by the earlywarnings R toolbox by Dakos et al [14]. (Parameters for simulation are as follows: *k*_offon_ = 0.043, *k*_onoff_ = 0.171, *k_p_* = 0, *k*_1_ = 2, *V^+^* ϵ [0,1000].)

### 3.2.2 Window size and white noise related sensitivity

By comparing figures S5, S6, and S7, we find that spike rate and spike span time is more robust, maintaining their sensitivity to the critical threshold while being less affected by the rolling window size and the filter window size. Figure S5 shows that an appropriate rolling window size and filtering window size is required for ar(1) to correctly indicate the transition. For a large rolling window, such as *η =* 0.31, ar(1) fails to predict the critical transition. A small filtering window also affects ar(1) performance by introducing noise. Figures S6 and S7 show that spike rates and spike span time indicate the critical threshold for any rolling window size *η >* 0.18, by showing a strong trend or peak near the threshold, and the filtering window has little effect on its performance. Spike rate and spike span time fails when a tiny rolling window *η =* 0.01 and a relatively large filtering window of *ζ >* 34 data points are used.

For extraneous white noise, parameter *c* is assigned 5 values within the range [10^3^, 10^7^] (section 2.4.2). After adding scaled noise, we perform indicator calculations. The results are shown in figure S8. Without additional noise, autocorrelation-at-lag-1 ar(1) and standard deviation (SD) are able to detect the critical threshold with a strong linear trend. With more noise, the trends in ar(1) and SD decrease until these indicators are no longer sensitive to the transition. From figure S8, when *c* ϵ [4×10^5^, 6×10^5^], the spike rates and spike span time are still sensitive to the threshold, showing peaking near the critical threshold, while ar(1) and SD trends toward transition is significantly decreased. Thus we conclude the spike rate and spike span time are more robust indicators compared to ar(1) and SD in our study.

### 3.2.3 Indicator significance

We performed the BDS test on the 8 different indicators shown in figure 4, including spike rate, spike span time, autocorrelation-at-lag-1, autoregressive-coefficient-at-lag-1, return rate, density ratio, SD, and skewness with the same simulated time trajectory for metabolite quantity, and found all *p*-values to be less than 0.0001. Thus we reject the i.i.d. null hypothesis and conclude that nonlinear structure is significant in the time series. Together with strong indicators shown in figure 4, the detected early warning is unlikely to be a false positive.

To assess early warning indicators, we propose a test model using a two-sample Z-test. The method is intended to determine if the difference in spike rates is significant between different system states. We simulated models with 20 linearly spaced *V^+^* ϵ [0,1000], with 100 repeats for each simulation. We tested the null hypothesis that there is no difference between spike rates of different system states. After obtaining mean spike rate and standard deviation for the 100 repeats, we performed the two-sample Z-test between pairs of conditions and ploted the p-values in figure S9b. Percentage enzyme saturation *P_es_* and percentage of trajectories that cross transition and their spike rates are shown in figure S9a; peak spike rates indicate the threshold. In figure S9b, region I tests the spike rate difference between post-transition (enzyme saturated) states and transitional states. Region II tests the spike rate difference between pretransition states (unsaturated enzyme) and the transitional state. Both region I and region II have small *p*- values less than 0.0001, thus rejecting the null hypothesis and leading to conclude that there is a significant difference in spike rate for different states. Other regions of the plot have *p*-values ~1, thus we fail to reject the null hypothesis when comparing the spike rates of pre-transition states and post-transition states, and conclude that there is no significant difference between spike rates of pre-transition states and post-transition states. In this way, with given information on stationary states, we can use a two-sample Z-test with sample spike rate data to predict an imminent transition. We can also apply this method with other early warning signals, such as spike span time.

### 3.2.4. Effects of Product Inhibition on System Dynamics

Product inhibition is a common metabolic phenomenon, commonly via allosteric or competitive inhibition of an upstream enzyme, and captures general effects of feedback on metabolic dynamics. To determine how negative feedback affects dynamical indicator behaviors, we considered metabolic product inhibition as shown in figure 2a (dashed arrows). In figure S11, the heatmaps show that different values of *k_p_* (the rate of metabolic product binding to enzyme) have different impacts on system dynamics. In the moderate feedback case, *k_p_ =* 10^-4^ (#s)^-1^, the system exhibits distinct saturation states with increasing *V*^+^ (figure S11a). In a heavy feedback case, *k_p_ =* 10^-1^ (#s)^-1^, there is a sharp change from unsaturated to saturated states. We found that both spike rate and spike span time are sensitive to system state transitions. While spike rates indicate the middle of the transition, spike span time indicates the end of the transition. Based on these benchmark simulation results, we chose *k_p_* values of 0, 10^-4^,10^-1^ (#s)^-1^ as representative in other parameter sweeps (indicated with light green line in figure S11).

### 3.2.5. Low Gene Burstiness Enhances Flickering

While molecular flickering in bacterial metabolic networks is likely to be a good indicator for phenotypic transitions, the generality of flickering as a predictor is still in doubt. Dakos *et al* studied dynamical system behavior with a simple model of additive tunable white noise, and concluded that trends in leading indicators (flickering) depend on the way noise affects dynamics in a system [15]. In metabolic networks, enzyme gene burstiness may affect phenotypic predictability. We thus carried out a controlled systemic comparison with simulations: using the same average enzyme expression level, we varied the gene burstiness, hence varying the shape of the enzyme distribution.

With our model, the average probability for a single copy gene to be on, *P_on_,* is given by
equation (5) and enzyme steady state given by
equation (10). As described in methods (section 2.4), changing *k_offon_* and *k_onoff_* proportionally will not change the *P_on_* value, and the steady state enzyme quantity in the system will stay the same.

We performed a set of simplified stochastic simulations with varying parameters to characterize the relation between the gene burstiness, *k_off_*_*on*_, and *k_onoff_*. In these simple simulations, only two reactions were considered: a single copy of the OFF-state gene switching between turning on and turning off, with respective propensities of: *P_off_* → *P_on_: k_offon_P_off_; P_on_* → *P_off_ : k_onoff_P*_*on*_. Parameter *P_on_* is assigned with 10 values spaced evenly in a linear space, starting from 10^-5^ to (1 — 10^-5^), and parameter *k_offon_* is assigned 80 values spaced evenly in a linear space starting from 10^-5^ to 2.0. The simplified model was simulated for 200 seconds with 200 repeats for each set of parameters (table S1). We quantify the gene burstiness by the frequency of enzyme gene expression changing states:

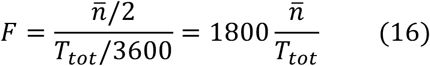

where *F* is the gene switch frequency, 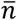 is the average gene state change counts for 200 repeats, and *T_tot_* is the total simulation time. *F* is therefore the average count of gene state changes per hour. We observed a non-linear increase in *F* with proportionally increased *k*_offon_ and *k*_onoff_, where *P_on_* stays the same (figure S10).

In the detailed metabolic step model (figure 2a), fixing *P_on_ =* 0.2, and varying *k_offon_* from 10^-8^ to 1 *s*^-1^ and *k_onoff_* according to
equation (11), we performed parameter sweeps for *k_offon_* with increasing *V*^+^. Each set of parameters was simulated with 100 repeats. As the percentage of saturated enzymes is correlated with the dynamical system state (section 2.5), we plotted *P_es_* in figure 5a to show the distinct states in parameter space. We manually set a hard cutoff to separate the *P_es_* into four distinct phases for visualization purposes. When *P_es_* ϵ [0,0.3), the system is not saturated, as shown in dark blue. When *P_es_* ϵ[0.3,0.7), the system is in the middle of the state transition, where the enzyme is approximately half saturated, as shown in light blue. The light blue part corresponds to the highest value of spike rates shown in figure 5b. When *P_es_* ϵ [0.7,0.9), the system is almost saturated is near the high steady state, shown in pink, which corresponds to the highest values of spike span times shown in figure 5c. When the enzyme gene is rarely in the ON state, i.e. *k_offon_* ϵ [10^-8^,10^-6^) s^-1^, there are no spikes because the enzyme quantity is always low. When the gene is moderately bursty, i.e. *k_offon_* ϵ [10^-6^,10^-3^) *s*^-1^, though the system is noisy, spike rates and spike span times have the highest value in the transition zone between unsaturated and saturated states. When the gene switches frequently between ON and OFF states, i.e. *k_offon_* ϵ[10^-3^,1) *s*^-1^, the system dynamics are more uniform, indicating that frequent gene state switching does not influence indicator ability to predict an impending threshold, as spike rates and spike span time are uniformly present in the phenotypic transition zone. We also considered cases with moderate and strong product inhibition, and similar dynamics were observed (figures S12 and S13). The detailed reaction scheme and parameters are shown in table S2.

**Figure 5.**
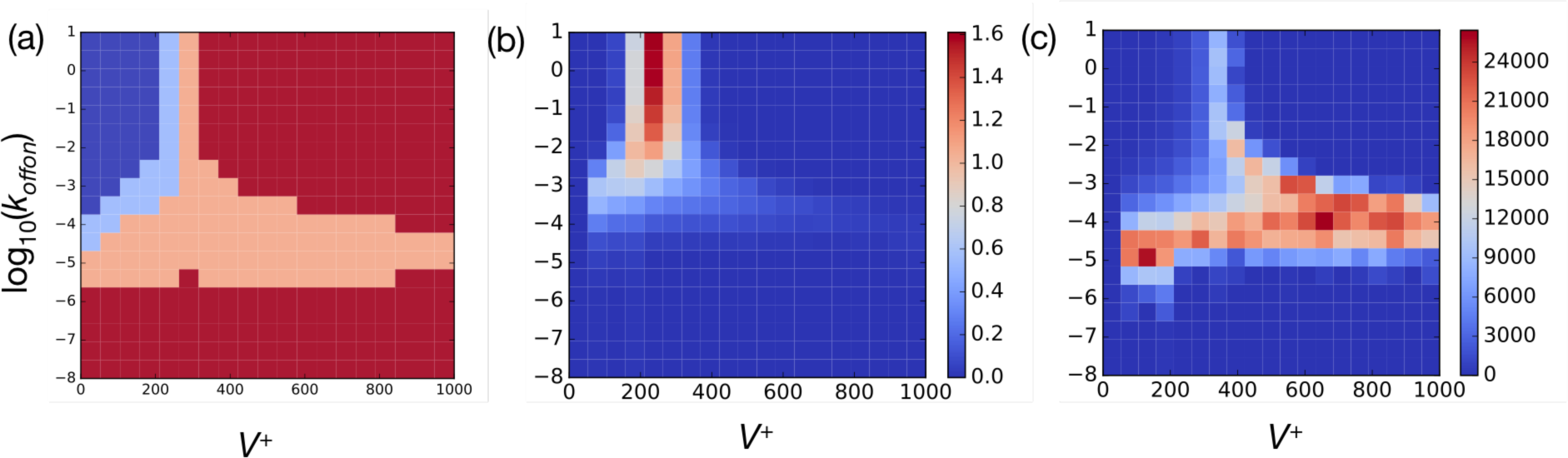
Heatplots of phenotype and indicators in the parameter space of gene burstiness and metabolic precursor influx *V*^+^ shows indicator dynamics. (a) System states in the simulation are shown with percentage of enzyme saturation. We set hard cutoff at *P_es_ =* 0.3, 0.7, 0.9 to visualize four distinct phases, and the two transition states in light blue and pink corresponding to the maximum of the spike rates and spike span time in panel (b) and (c). (b) Spike rate heatplot shows that the spike rates are sensitive to the middle of the transition. (c) Spike span time heatplot shows that the spike span time is sensitive to the near-saturation state of the system.

Overall, we found that flickering in regulatory networks can predict impending phenotypic transitions in enzymes with various levels of gene burstiness, except in cases where the threshold is so sensitive that no flickering can occur during transition processes. When gene burstiness is low enough, with an initial gene state of OFF, a lack of enzyme expression eliminates apparent flickering.

### 3.2.6. Dynamical effects of metabolic precursor binding affinity

With the idea that the spike rates and span time may predict the threshold transition, we further conducted parameter sweeps for the metabolic precursor binding rate *k*_1_ and metabolic precursor influx *V*^+^, and each set of parameters was simulated with 100 repeats. Metabolic precursor binding rate *k*_1_ is a measure for substrate binding affinity and inversely contributes to the classic Michaelis-Menten constant *K*_M_, which has a wide range of values for different enzymes:

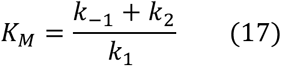

where *k*_-1_ is the metabolic precursor-unbinding rate and *k_2_* is the catalysis rate. The detailed reaction scheme and parameters are shown in table S3. The simulation results are shown in figure 6. From figure 6a, we observe that low substrate affinity with log(fc_x_) < −5 results in no saturation at all for any possible influx *V*^+^ value. Low substrate affinity also introduces noise to the system dynamics of the indicators, as we observe noisy spike rates in corresponding parameter space, while spike span time is less noisy. For log_10_(*k*_1_) ≥ −5, we observe distinct system saturation schemes that correspond to spike rates and spike span times in parameter space, similar to results discussed in the gene burstiness case. Metabolic product inhibition was also varied in this case, and the simulation results show similar traits, with negative feedback altering system dynamics, but with the spike rate and spike span time still able to indicate the threshold states (figures S14 and S15).

**Figure 6.**
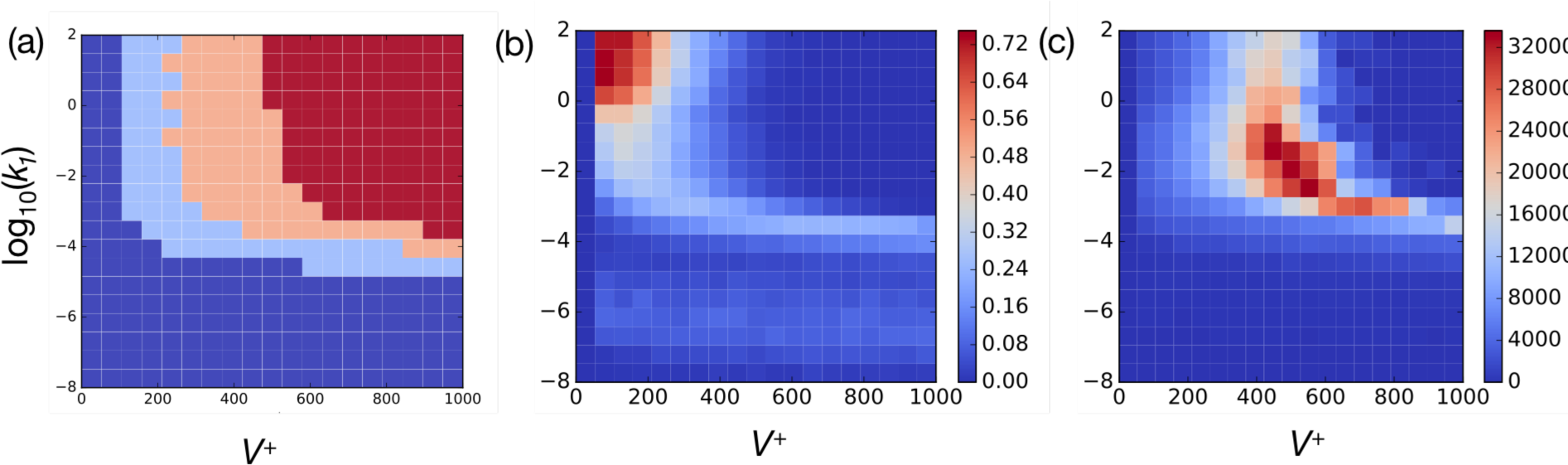
Heatplots for simulated system phase and corresponding indicator dynamics. We scanned the parameter space for metabolic precursor binding rate *k*_1_ and metabolic precursor influx *V*^+^ in the model with no feedback. (a) Phase graph shows non-saturation regime (deep blue), medium saturation regime (light blue), near-saturation regime (pink), and saturation regime (red) for different *P_es_* values. We manually set hard cutoff at *P_es_ =* 0.4,0.8,0.98 for visualization purposes. (b) Spike rate heatplots in parameter space shows that the spike rates correspond to the moderate saturation regime in (a) and indicate the system is crossing the saturation threshold. (c) Spike span time heatplot in parameter space shows that the spike span time, which can be regarded as the return time for the system, corresponds to the near-saturation regime, where the system is almost finished crossing the threshold.

Thus, simulations varying substrate affinity again yield the conclusion that spike rate can indicate the center of an ultrasensitive threshold transition, and spike span time can indicate the near-saturation states in a metabolic step. While low substrate affinity causes noise in system dynamics with concomitant increased spike rates, we consider this a false-positive case, detecting fluctuations of metabolite production and growth-mediated degradation. This noise is not obvious in the spike span time readout, further suggesting that these are not "true" spikes.

### 3.2.7. Dynamical Effects of Enzyme Gene Transcription Rates

In biological molecular regulatory networks, transcriptional noise is an important component of intrinsic noise [31], and phenotypic switches are often driven by transcriptional regulation [32, 33]. We therefore performed parameter sweeps to explore how enzyme expression level impacts fluctuations of metabolite (figure 7). As before, we considered three cases of metabolic product inhibition for the simulations: *k_p_=* 0,10^-4^,10^-1^ (#s)^-1^. The detailed reaction scheme and parameters are shown in table S4.

**Figure 7.**
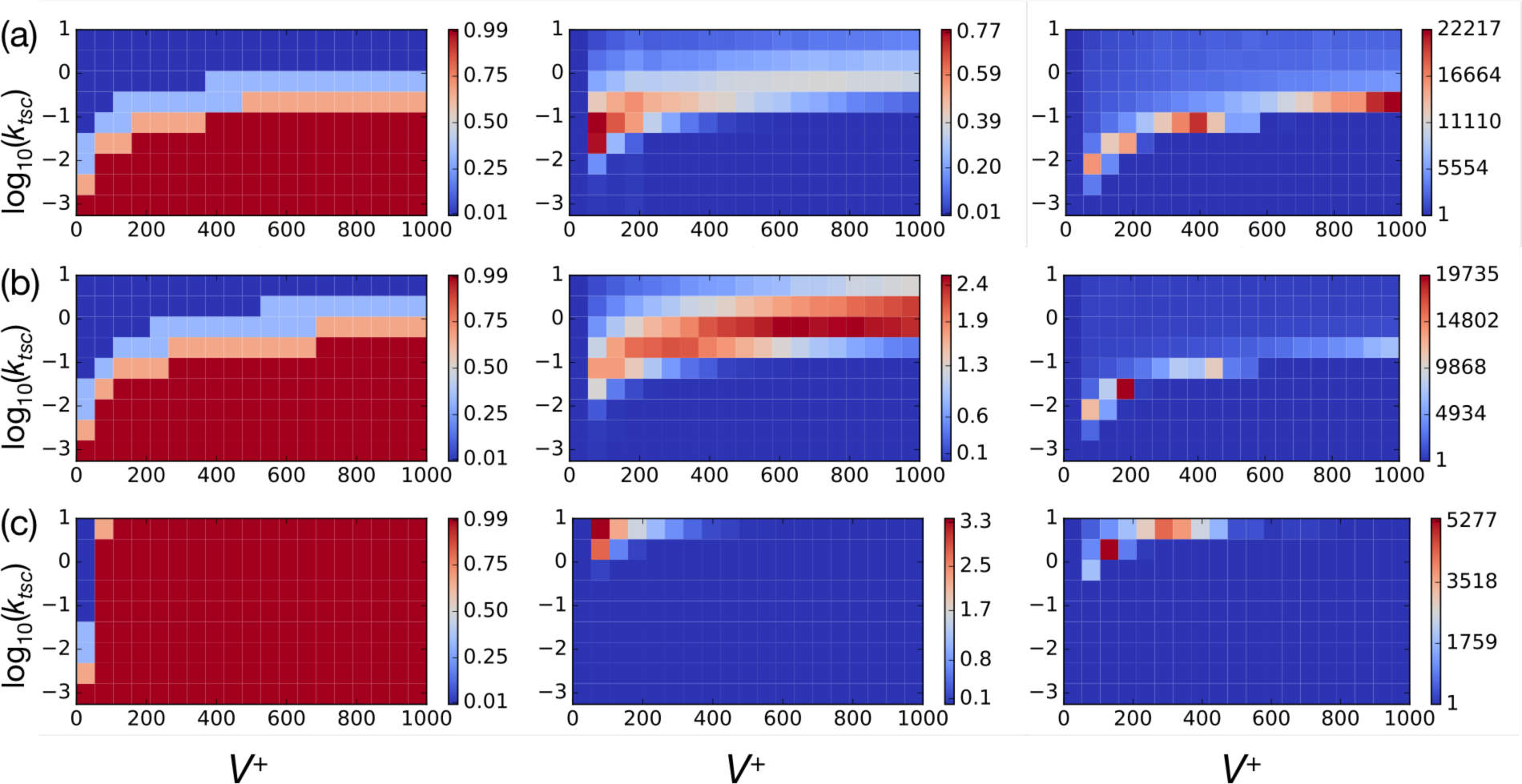
Heatmaps for simulated system phase and corresponding indicator dynamics with varying levels of product inhibition feedback. We scanned the parameter space of enzyme gene transcription rate *k_tsc_* and metabolic precursor influx *V*_+_. Here we present all three cases considering the metabolic product *P* competitive inhibition with product binding rate *k_p_ =* 0,10^-4^,10^-1^ (#s)^-1^. (a) Heatplots for no product inhibition case, where *k_p_ = 0* (#s)^-1^. For visualization purpose, we set a hard cutoff at *P_es_ =* 0.3,0.7,0.95 for the saturation phase graph (left panel); corresponding spike rates and spike span times are shown in the middle and right panel, respectively. Similar to (a), heatmaps (b) and (c) show system phase, corresponding spike rates and spike span time for moderate product inhibition *k_p_ =* 10^-4^ (#s)^-1^ and strong product inhibition *k_p_ =* 10^-1^ (#s)^-1^. Negative regulation of the pathway by product inhibition impacts system dynamics, while spike rates and spike span times are able to indicate the transition as long as the threshold is not too sensitive.

We observe different system dynamics depending on the strength of product inhibition. With higher product binding affinity *k_p_,* the thresholds between unsaturated states and saturated states arise at lower substrate concentrations, thus affording less parameter space for spikes to emerge. We also observe similar dynamics that spike rates indicate the intermediate saturation states, and spike span times indicate near-saturation states, which are consistent with previous simulations.

We found a higher maximum spike rate in feedback cases (figure 7b, 7c; figures S16 and S17). Metabolic precursor spiking originates from the timescale difference between faster metabolic fluctuation and the slower process of enzyme synthesis. From observation of exemplar cases, in a system with higher metabolic product binding affinity, enzymes tend to be more saturated, thus dragging the system state toward the saturation threshold (figure 8a, 8b). With higher *k_p_*, smaller *V*^+^ is required to saturate enzymes. We also scanned the parameter space for enzyme gene burstiness, substrate affinity, and transcription rates with different product inhibition strength, showing consistent results (figure S16).

**Figure 8.**
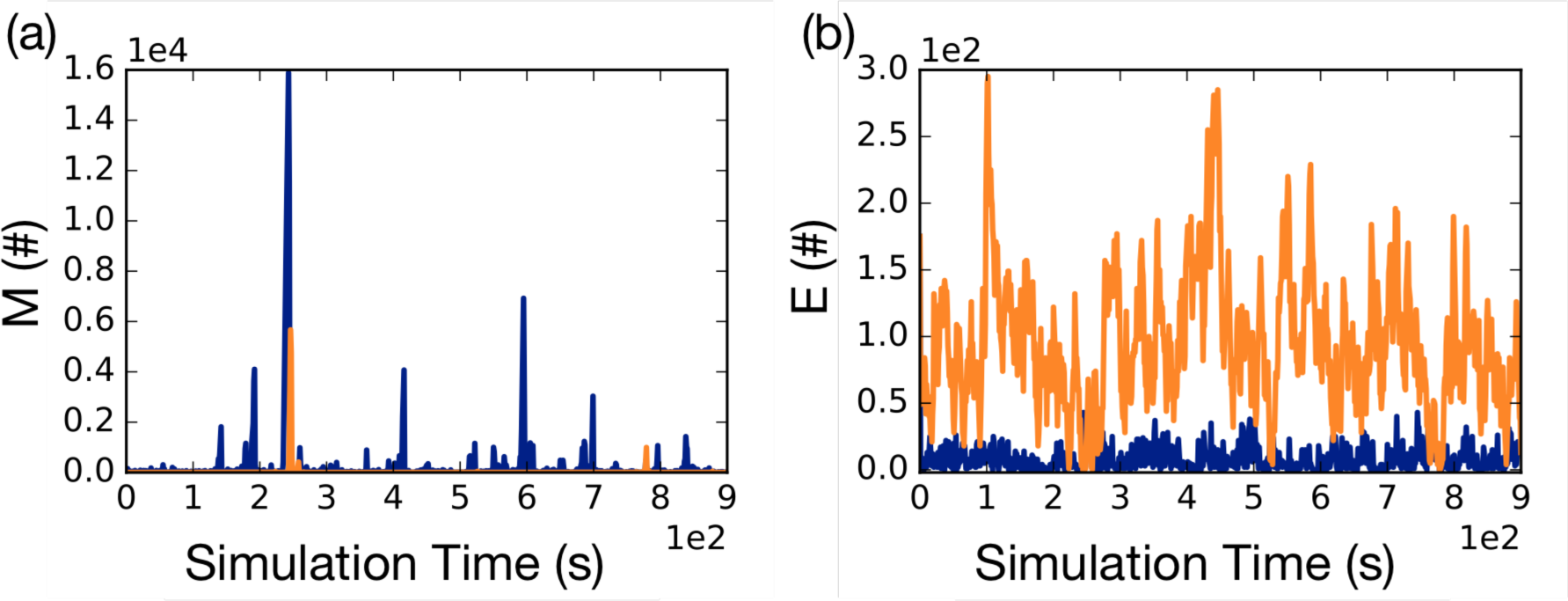
Higher spike rates are observed in negative feedback cases. Blue line indicates the time trajectories for higher metabolic product inhibition, where *k_p_=* 5.46×10^-4^ (#s)^-1^. Orange line indicates the time trajectories for lower metabolic product binding inhibition, where *k_p_=* 2.07 × 10^-5^ (#s)^-1^. (a) Metabolic precursor time trajectories are compared in two cases, and higher spike rates is observed in the stronger feedback case. (b) Free enzyme comparison shows that enzymes are more saturated (blue line) under stronger competitive inhibition from the product.

### 4. Discussion

We used a computational approach to study phenotypic switches caused by bacterial metabolic dynamics, and unveil spiking of metabolite (flickering) as an early warning indicator for predicting ultrasensitive thresholds in metabolism. With our simulation framework, we discovered that while parameters can significantly impact the ultrasensitive responses of enzyme saturation [34], flickering as an early warning indicator is able to predict the transition in a robust parameter space: as long as gene expression is not too bursty (figure 5) and background noise is not extreme (figure S8). Our minimalistic models achieve distinct phenotypic regimes through varying rates of metabolite precursor influx, enzyme gene burstiness, expression level, metabolic precursor binding rates, and feedback from product inhibition. We demonstrate that peaks of flickering rates and spike span time correspond to the switching process wherever the saturation threshold is not too low, thus providing a potential pathway toward developing methods for predicting phenotypic transitions in single bacterial cells by tracking the quantity of molecules in natural regulatory networks or engineered synthetic circuits.

This effort is motivated by the field of early warning indicator studies for predicting catastrophic transitions in complex networks. Considering multiple similarities between cellular regulatory networks and complex systems in ecology and economics, including many interactions, intricate network structures, and ultrasensitive thresholds, we seek similar methods for predicting critical transitions in single biological cells. With this study, we discovered consistent dynamical behaviors in metabolic steps as rising noise and increased return time, comparable to other complex systems. We found increasing deviation near the enzyme saturation threshold in simulations and analysis, characterized spike rate and long spike span time as early warning indicators in our system, and demonstrated the usefulness of other established indicators. Despite this similarity, there are differences in the function of flickering between the metabolic step and most other complex systems analyzed with early warning indicators: flickering in our study arose not from fluctuations between different steady states, but rather, from the shift between different potential regimes at the critical point (figure 3c). Thus, there is only one steady state for each set of parameters in the deterministic model, while noise in the system may frequently push the system to switch potential regimes near the threshold.

Our method shows that flickering in regulatory networks can be used as an early warning indicator for a large segment of parameter space, whenever the threshold is not too sensitive. It is worth noticing that spike rates indicate the state in the middle of a transition, while spike span times indicate a near-saturation state. Longer spike span times may relate to the phenomenon of critical slowing down that is often observed at such thresholds in dynamical systems. Apparent false-positive cases can arise when we have extremely low enzyme affinities, effectively having a system of Poissonian production and degradation of metabolite without significant consumption by enzyme. Because spike span times indicate a near-saturation regime, it behaves better than spike rates against false-positive indication. Both indicators behave for many different parameter combinations, suggesting that flickering dynamics are not parameter-directed, but rather system state-directed, which is promising for its generality in applying to different biological systems.

Our study demonstrates that early warning indicators can be useful in cellular regulatory networks. With our methods, we would be able to tell the approximate system states by observing fluctuations of certain compounds in single cell with a set of data. This would be helpful for evaluating cell status when establishing animal disease models in a medical research. With help of time-lapse microscopy, we should be able to predict single cell phenotypic transitions if the observed indicator is a visible molecule, such as a fluorescent protein. To be useful for detecting critical transitions, the indicator molecule must have detectable fluctuations with a high enough rate that indicators can be reliably computed. It thus may help us further understand the underlying mechanism of cell fate decisions and enable markers to intervene with this process.

It is additionally possible that fast fluctuations may somehow signal an imminent transition to other components of the regulatory network. Gene expression or other regulatory network states may then be poised for a transition that has not yet occurred, signaled by the fluctuations of an indicator molecule.

### 5. Conclusions

Within a reasonable range of parameters, one-enzyme metabolic steps display early warning indicators of being close to the critical transition between saturated and unsaturated enzyme states. Additional work is needed to determine the generality of such indicators in larger metabolic networks and in other regulatory networks containing processes with diverse timescales. We propose that dynamical early warning indicators could be utilized in clinical applications or synthetic biology as a readout of cells near a critical phenotypic transition point, and that such fluctuations may affect the physiology of cells that are near the critical point.

## Acknowledgments

We thank Will Mather for helpful comments. This project was supported by the National Institute of General Medical Sciences (P20 GM103418) from the National Institutes of Health. The content is solely the responsibility of the authors and does not necessarily represent the official views of the National Institute of General Medical Sciences or the National Institutes of Health.

